# Genetic and transcriptional analysis of human host response to healthy gut microbiome

**DOI:** 10.1101/058784

**Authors:** Allison L. Richards, Michael B. Burns, Adnan Alazizi, Luis B. Barreiro, Roger Pique-Regi, Ran Blekhman, Francesca Luca

**Affiliations:** 1Center for Molecular Medicine and Genetics, Wayne State University, Detroit, MI; Departments of Genetics, Cell Biology, and Development, The University of Minnesota, Minneapolis, MN; Department of Ecology, Evolution, and Behavior, The University of Minnesota, Minneapolis, MN; Department of Pediatrics, Sainte-Justine Hospital Research Centre, University of Montreal, Montreal, QC, Canada; Department of Obstetrics and Gynecology, Wayne State University, Detroit, MI

## Abstract

Many studies have demonstrated the importance of the gut microbiome in healthy and disease states. However, establishing the causality of host-microbiome interactions in humans is still challenging. Here, we describe a novel experimental system to define the transcriptional response induced by the microbiome in human cells and to shed light on the molecular mechanisms underlying host-gut microbiome interactions. In primary human colonic epithelial cells, we identified over 6,000 genes that change expression at various time points following co-culturing with the gut microbiome of a healthy individual. The differentially expressed genes are enriched for genes associated with several microbiome-related diseases, such as obesity and colorectal cancer. In addition, our experimental system allowed us to identify 87 host SNPs that show allele-specific expression in 69 genes. Furthermore, for 12 SNPs in 12 different genes, allele-specific expression is conditional on the exposure to the microbiome. Of these 12 genes, eight have been associated with diseases linked to the gut microbiome, specifically colorectal cancer, obesity and type 2 diabetes. Our study demonstrates a scalable approach to study host-gut microbiome interactions and can be used to identify putative mechanisms for the interplay between host genetics and microbiome in health and disease.

## Importance

Study of host-microbiome interactions in humans is largely limited to identifying associations between microbial communities and host phenotypes. While these studies have generated important insight on the link between the microbiome and human disease, assessing cause and effect relationships has been challenging. Although this relationship can be studied in germ-free mice, this system is costly, and it is difficult to accurately account for the effect of host genotypic variation and environmental effects seen in humans. Here, we have developed a novel approach to directly investigate the transcriptional changes induced by live microbial communities on human colonic epithelial cells and how these changes are modulated by host genotype. This method is easily scalable to large numbers of host genetic backgrounds and diverse microbiomes, and can be utilized to elucidate the mechanism of host-microbiome interactions.

## Introduction

A healthy, human adult contains over one thousand species of bacteria in their gut (1). These bacteria live in a symbiotic relationship with us and compose the gut microbiome. Recent studies suggest that the gut microbiome may play a role in both physiological and pathological states. The composition of the gut microbiome has been correlated with complex diseases, such as Crohn’s disease and diabetes (2–5). The two most abundant phyla in the human gut are bacteroidetes and firmicutes (1). In obese individuals, the ratio of these two phyla is altered (6–8). Turnbaugh et al. showed that transplanting the fecal microbiome of an obese mouse to a germ-free mouse causes greater weight gain in the recipient as compared to recipients that received the microbiome of lean mice (9). Goodrich et al. showed that this relationship exists even when the microbiome from obese humans is transplanted into mice (10). The microbiome has also been linked to colorectal cancer (11, 12) and to diseases not directly related to the gut, such as arthritis, Parkinson's disease, and other types of cancer (13–16).

While there are many species that are common among humans, studies have shown that microbiome composition can vary widely across individuals (17, 18). These differences have been correlated to several factors, such as breastfeeding, sex, and diet (19–24). In addition to environmental factors, recent studies also support a key role for host genetics in shaping the gut microbiome. Indeed, microbiome composition is more similar in related individuals than in unrelated individuals (10, 25–28). One caveat of these studies is that, especially in humans, related individuals often share environments and follow similar eating habits, which have a strong effect on the microbiome. In an effort to control for this factor, other studies have attempted to estimate the role of host genetics on the microbiome in mice, where the environment can be regulated, or in groups of people that all share the same environment regardless of relatedness (29–32).

To further examine the effect of host genetic variation on gut microbiome, some groups have performed association studies between host genotypes and microbiome composition (32–35). For example, Blekhman et al. studied 93 individuals and identified loci that are associated with microbiome composition in 15 body sites that were sequenced as part of the Human Microbiome Project (18, 33). Among SNPs associated with microbiome composition, they found an enrichment in SNPs that were identified as expression QTLs (eQTLs) across multiple tissues in the Genotype-Tissue Expression (GTEx) project (36). Additionally, microbiome composition has been found to be tissue-specific and therefore, likely influenced by host gene expression pattern in the specific tissue that interacts with the microbiome. Together, these results suggest that host genetic variants affects microbiome composition through influencing host gene and protein expression. However, we know little about the interplay between human genetic variation, gene expression, and variation in microbiome composition, and the effect of these factors on susceptibility to complex disease.

Molecular studies of genetic effects on cellular phenotypes (eQTL, dsQTL and transcription factors binding QTL mapping studies) have been successful in elucidating the link between genetic variation and gene regulation, and have identified hundreds of variants associated with gene expression and transcription factor binding changes (37–42). Here, we present a novel approach to study the interaction between the microbiome, human genetic variation and gene expression in a dynamic and scalable system. We co-cultured primary, human colonocytes (epithelial cells of the colon) with the gut microbiome of a healthy individual (extracted from a fecal sample) to study host cell gene expression response to microbiome exposure. We identified over 6,000 genes that significantly change their expression in the host following microbiome exposure. These genes are enriched for GWAS signals, suggesting that regulation of their expression is a potential mechanism for the associations found between host disease status and microbiome composition. In addition, to learn about host genetic variants that play a role in host-microbiome interaction, we studied allele-specific expression (ASE) and identified 12 genes that demonstrate an interaction between genotype and microbiome exposure. Future studies can use this approach to characterize host response to the microbiome and determine the causal relationship in the context of specific diseases and traits.

## Results

### Study design

While many recent studies have shown the importance of the microbiome in physiological and pathological states, in humans, the direct impact of exposure to the microbiome on host cells is yet unclear. To analyze the host transcriptional changes induced by a normal gut microbiome, we designed an experiment in which we co-cultured human colonic epithelial cells (colonocytes) with an extract containing the fecal microbiome from a healthy individual (**Figure 1A** and Table S1). We analyzed the DNA of the fecal extract through 16S sequencing followed by data processing using QIIME (12, 43, 44) to quantify microbial species present. This fecal extract showed a normal composition of bacteria phyla with Firmicutes and Bacteroidetes representing the most abundant taxa, consistent with previous studies of gut microbiota composition in healthy individuals (Table S1) (18).

**Figure 1:**
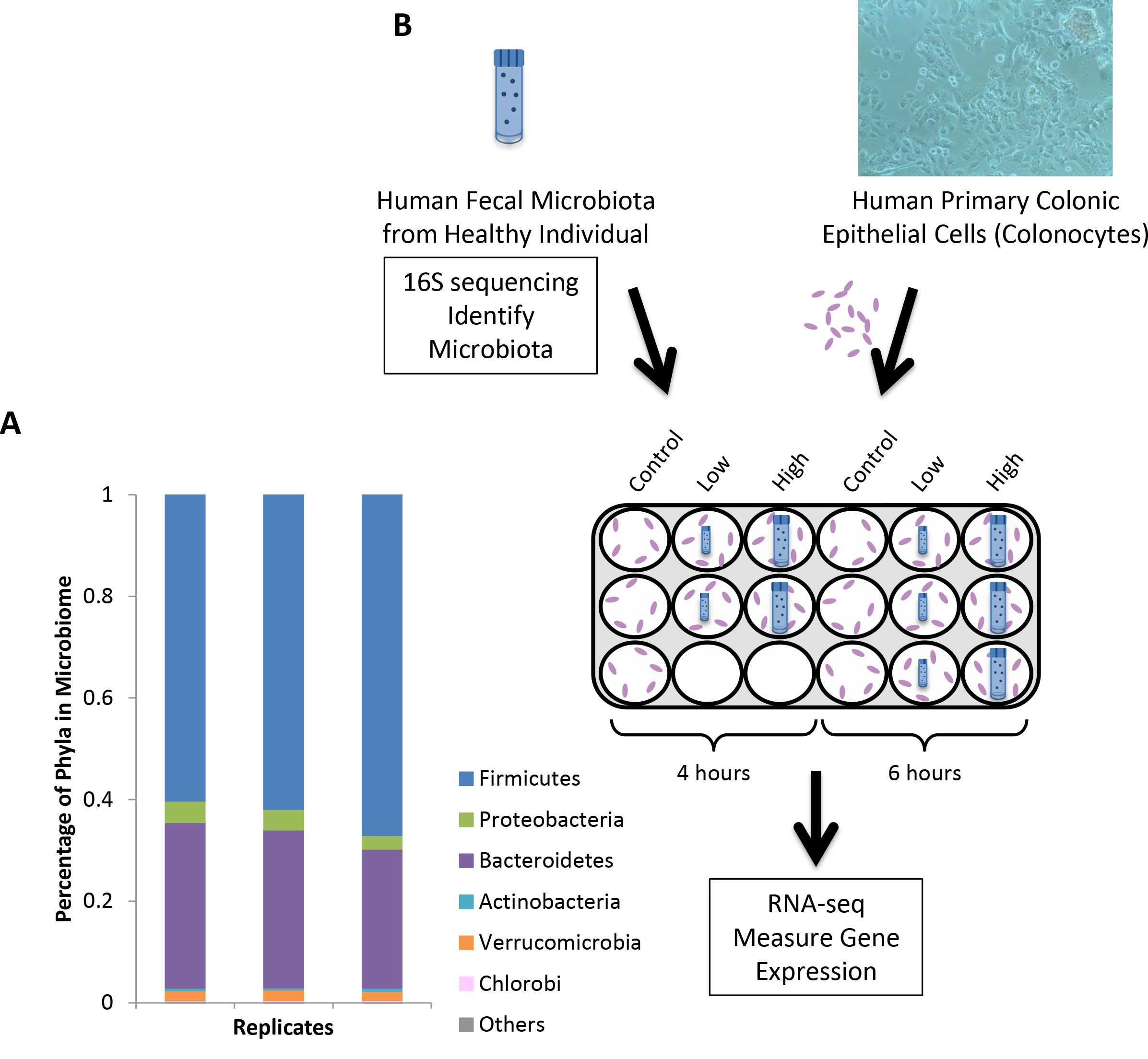
Co-culturing of human colonocytes and fecal extract. A) 16S rRNA gene sequencing results from fecal extract of healthy, 22 year old male used to co-culture with colonocytes. Each bar denotes a replicate of the same uncultured fecal extract. The most abundant phyla are depicted as a percentage of the total microbiome detected. B) Treatment scheme to co-culture colonocytes and microbiome which was then followed by RNA-sequencing of mRNA to assess host gene expression. Cells were treated for 4 and 6 hours using a high or low concentration of fecal extract (or no fecal extract as controls).

We exposed the colonocytes to two different densities of live microbiome extract (as measured by 0D600), including 10:1 and 100:1 bacteria:colonocyte ratios, termed High and Low concentration, respectively. We cultured the colonocytes in low Oxygen (5% O_2_) to recapitulate the gut environment for 4 and 6 hours under three conditions: with high and low concentrations of bacteria and alone, as controls (**Figure 1B**). This resulted in 5 experimental conditions: Low-4, Low-6, High-4, CO4 and CO6. Experimental replicates were collected for each condition: two replicates for Low-4 and High-4 and three replicates for Low-6, CO4 and CO6. We collected and sequenced the RNA in order to learn about the host cell response through study of gene expression and to identify genes with allele-specific expression induced by the microbiome.

### Transcriptional changes induced by the gut microbiome

First, we searched for genes that were differentially expressed (DE) in the colonocytes following exposure to the gut microbiome. We used DESeq2 (45) as described in the methods to characterize differential gene expression in the treatment samples, across biological replicates. We focused on genes with significant differences using a Benjamini-Hochberg adjusted p-value < 0.1 and |log_2_(Fold-Change)| > 0.25. With this method we identified 3,320 genes that change expression in Low-4 relative to CO4 (55% up-regulated), 1,790 genes in Low-6 relative to CO6 (57% up-regulated) and 5,182 genes in High-4 relative to CO4 (49% up-regulated) resulting in 6,684 genes that had at least one transcript that was differentially expressed (DE) under any of the three conditions (**Figure 2**, Figure S1, and Table S2).

**Figure 2:**
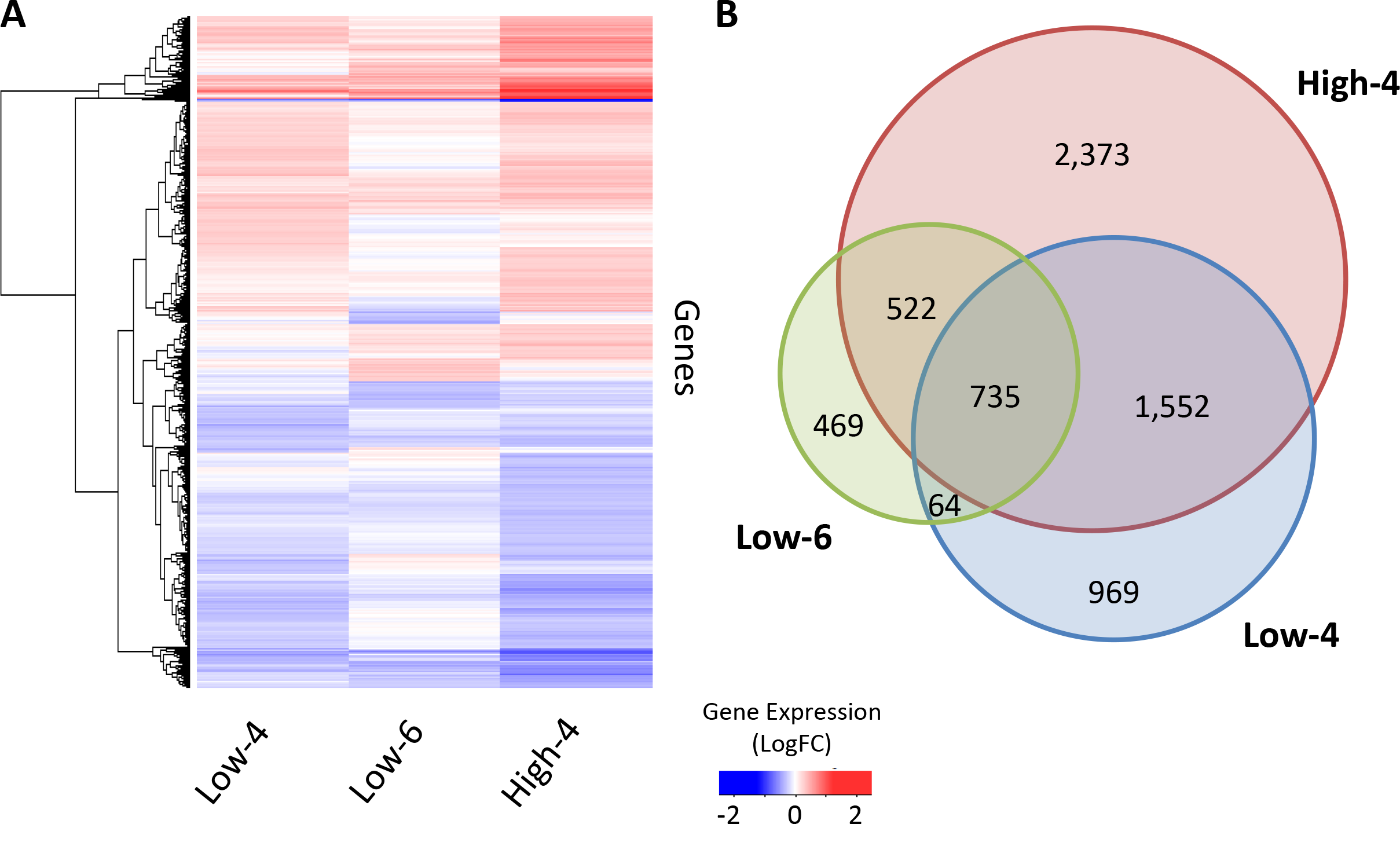
Host gene expression changes following exposure to the microbiome. A) Heatmap depicts average across replicates of log_2_(Fold Change) for each sample as compared to the respective control (Low-4 and High-4 are compared to CO4 while Low-6 is compared to CO6). Blue indicates a decrease in expression in the treatment sample while red indicates an increase in expression. One transcript from each of 6,684 genes that are DE in any of the 3 treatments (Benjamini-Hochberg adjusted p-value < 0.1, |logFC| > 0.25) is shown. B) The Venn diagram depicts the number of genes that contain any transcript differentially expressed under the various treatment conditions. The overlap numbers require that the same gene is DE in the different samples.

To determine whether our results recapitulate gene expression patterns observed in *in vivo* models, we performed a comparison to an existing dataset assessing the effect of the microbiome on colonic gene expression in mouse (46). Camp et al. studied mice that were in three groups: conventionally raised (CR), mice raised in a germ-free environment that were then conventionalized with microbiome for 2 weeks (CV) and mice only raised in a germ-free environment (GF) (46). They performed RNA-seq and identified 194 and 205 genes that were differentially expressed in colonic epithelial cells in CR and CV mice, respectively, as compared to GF mice. When we searched for the overlap between our 6,684 differentially expressed genes we found that we had a significant enrichment for the genes differentially expressed in CR mice (42 genes out of 194 DE genes in CR mice, Fisher's Exact test p-value = 0.001, OR = 2.3) but not with CV mice (Fisher's Exact test p-value = 0.39). This suggests that our model more accurately represents a normal, healthy interaction with the microbiome as compared to the acute response observed in the CV mice.

We next examined the function of the genes that change their expression in the host. We identified genes involved in pathways previously shown to be affected by exposure to microbiome, including cell-cell junctions (47, 48) and lipid metabolism (46, 49) (**Figure 3A**). Similar to Camp et al. (46), we also identified changes in gene expression of transcription factors. Specifically, we find an enrichment of genes for which Camp et al. found binding sites near genes differentially expressed between CV and CR mice (50 transcription factors that are DE in our data) (Fisher’s exact test p-value = 0.0004, OR = 2.3). This includes *EGR1*, a gene involved in intestinal response to injury (50), and several *STAT* genes, which are part of a pathway up-regulated in colorectal cancer (51). This overlap suggests that our *in vitro* system accurately depicted an *in vivo* response and that the changes in host gene expression are mediated by changes in the abundance of key transcription factors in humans as Camp et al. had seen in mice.

**Figure 3:**
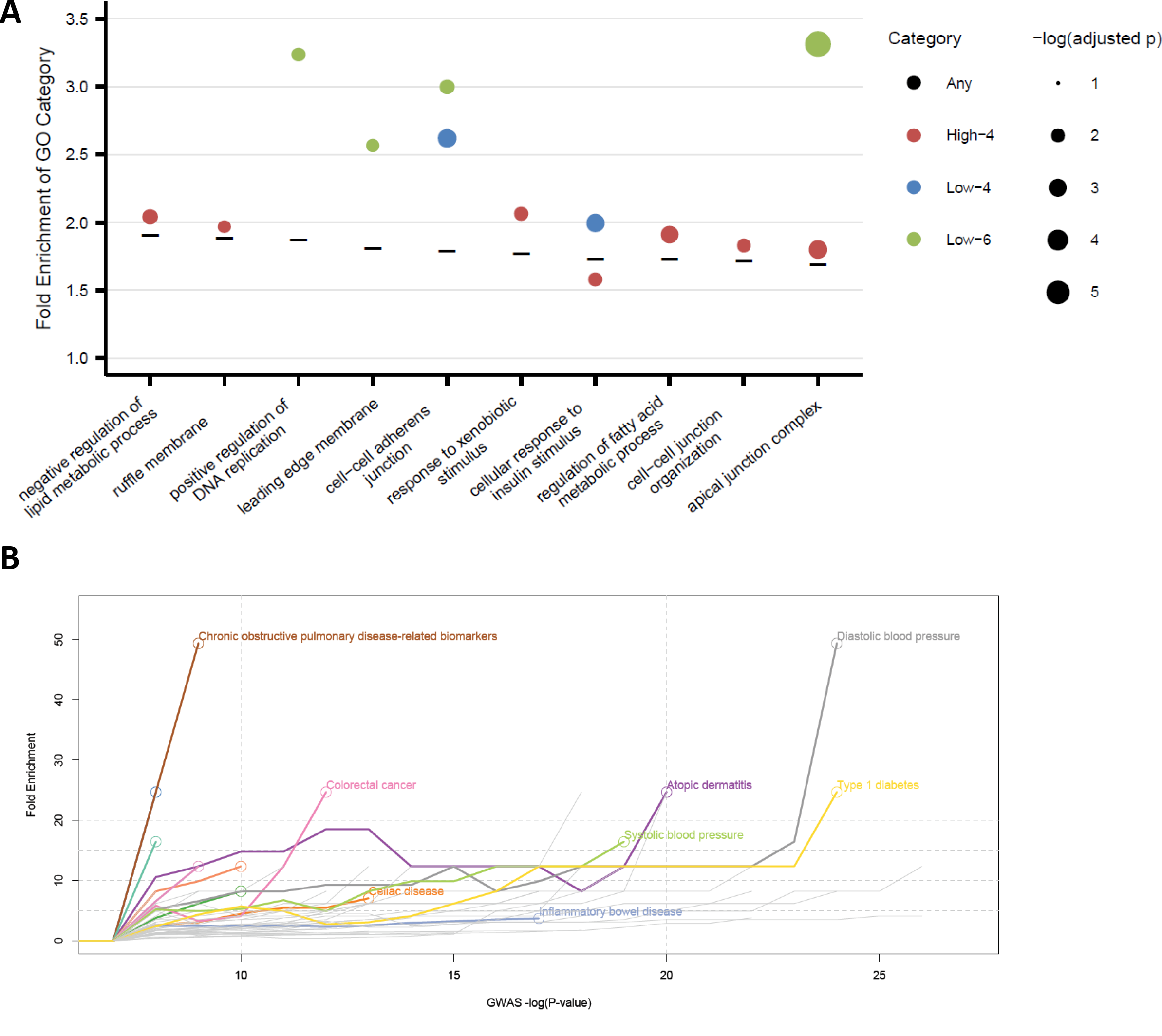
Functional enrichment of differentially expressed genes. A) GO enrichment was assessed using GeneTrail (87) for any gene differentially expressed in any of the 3 treatments (6,684 genes). Enrichments for the top 10 categories that were over-represented are indicated with a black bar (details in methods). GO enrichment was performed for genes differentially expressed in each of the 3 treatments separately and if these categories were significantly overrepresented, the enrichment in that category is shown by a closed circle (Low-4 is blue, High-4 is red, Low-6 is green). The closed circles are weighted based on the -log_10_(Benjamini-Hochberg adjusted p-value). C) Fold enrichment of DE genes (y-axis) among genes associated in GWAS for a given disease at progressively stringent p-value thresholds (x-axis). For each GWAS and P-value cutoff, we identified the overlap between the genes significantly associated with the disease at that cutoff and DE genes in our study, and calculated a fold enrichment (plotted along the y-axis), defined as the ratio of observed/expected overlap between the two gene sets. Colored lines indicate an enrichment significant at p < 0.05 (using a Fisher’s exact test), with the point of maximum enrichment indicated by a circle. The GWAS disease name is listed next to the line for diseases with a fold enrichment > 30 or x-axis position of maximum enrichment > 10.

Previous reports in animal models have demonstrated enrichment for genes involved in immune response among those that change expression following short-term and long-term exposure to the microbiome (46, 52, 53). Indeed, among the GO categories that are significantly enriched (Benjamini-Hochberg adjusted p-value < 0.05) with genes that change expression following co-culturing we find immune system process. We wondered whether immune response activation is stronger under certain conditions. Specifically, we tested whether the high dose of microbiome at 4 hours had a stronger effect on the immune response than the low dose at 4 hours. We identified 2,094 genes that were differentially expressed between High-4 and Low-4, with transcripts from 1,308 genes showing increased expression at the higher concentration and transcripts from 788 genes showing decreased expression at the higher concentration of microbiome (Table S3). When we searched among the genes that are increased in expression with the higher concentration of microbiome, we found several immune-related GO categories (Figure S2). These data suggest that a higher microbiome concentration elicits a stronger immune response in host cells.

### Transcriptional response and human diseases

The impact of microbiome exposure on gene expression led us to ask whether these changes may affect human diseases. Several diseases have been linked to variation in the composition of the gut microbiome, including obesity, type 2 diabetes, inflammatory bowel disease, Crohn’s disease, ulcerative colitis, and colon cancer (7, 27, 54–61). Many GWAS studies have identified genetic loci associated with these diseases (62), but in most cases, the mechanism by which the gene influences the disease is still unclear. Similarly, the mechanisms by which microbiome composition may influence human diseases are mostly still unknown. Our data allowed us to investigate these questions using primary human colonic cells.

First, we hypothesized that if we identify a differentially expressed gene in our data that is also associated with a disease, it is likely that changes in the microbiome influence the gene’s expression, thereby contributing to the health of the host. To test this hypothesis, we studied genes that were previously reported to be associated with any complex trait (NHGRI GWAS database) (62), as defined in the Methods. We searched among genes that were differentially expressed, in the same direction, in all 3 treatments and found enrichment for genes associated with complex traits (Fisher’s Exact test p-value < 10^−10^, OR = 1.8). We then focused on several diseases that have already been linked to microbiome composition. We found that DE genes were enriched for genes associated with obesity-related traits (Fisher’s Exact test p-value = 0.03, OR = 1.5) and colorectal cancer (Fisher’s Exact test p-value = 0.01, OR = 3.0) with suggestive enrichment for inflammatory bowel disease (Fisher’s Exact test p-value = 0.06, OR = 1.7) and ulcerative colitis (Fisher’s Exact test p-value = 0.09, OR = 1.9). There was not significant enrichment for type 2 diabetes or Crohn’s disease (Table S4). Additionally, we found that the enrichment of genes associated with colorectal cancer is significant also when we used a complementary approach that accounts for the differences in the distribution of p-values across GWAS (**Figure 3C**). For this analysis, we used a range of −log_10_(p-value) cut-offs for each disease in the GWAS catalog, and identified the overlap between the genes significantly associated with the disease at each cutoff and DE genes in the current study. Using this approach, we also found enrichment among several autoimmune diseases that have been previously linked to variation in the microbiome, such as atopic dermatitis, celiac disease, and inflammatory bowel disease (63–65). These results support our system as a useful method for studying genes and interactions involved in organismal traits.

Moreover, dysregulation of the genes that are both differentially expressed and associated with these diseases may represent a mechanism that causes the pathological state through host cell response to the gut microbiome. Future studies utilizing microbiomes from healthy and diseased individuals will be able to further shed light on how different microbes may influence disease risk through changes in host gene expression.

### Allele-specific expression

Genetic variants associated with microbiome composition have previously been linked to expression changes in humans through eQTL studies (33). However, to date, there are no reports in humans on the effects of genetic variants on the host transcriptional response to the microbiome. In order to identify genetic loci that may influence host gut-microbiome interactions through their influence on gene expression, we studied allele-specific expression (ASE) (37–42). This analysis is ideal for our study (using colonocytes from a single individual) as it uses the genotypes and allelic imbalance for each individual separately to assess genetic control, as opposed to using multiple individuals to determine a correlation in a population between genotypes and expression (37–42). The caveat is that we can only assess SNPs that are heterozygous in our sample and deeply covered by sequencing reads. To characterize ASE in our samples, we utilized QuASAR (66), a method to detect heterozygous sites in a sample and utilize these sites to identify ASE. We found an average of 5,984 heterozygous SNPs per sample covered by at least 20 RNA-seq reads. Among these heterozygous sites, we identified 131 events of ASE at 87 SNPs in 69 unique genes (Storey FDR < 10%) across our samples, including controls (**Figure 4** and Tables S5 and S6). Three of these SNPs show the same ASE in all samples suggesting that these may play a role in the baseline regulation of colonocytes. 40 ASE events occur in the treatment samples and 18 occur in genes that are differentially expressed at the same time point. This suggests that these ASE events may be a result of either new transcription of the favored allele or specific degradation of the other allele. The 22 remaining ASE events may involve genes where there are changes in expression of transcripts containing both alleles such that the gene expression remains constant though the ASE may change.

**Figure 4:**
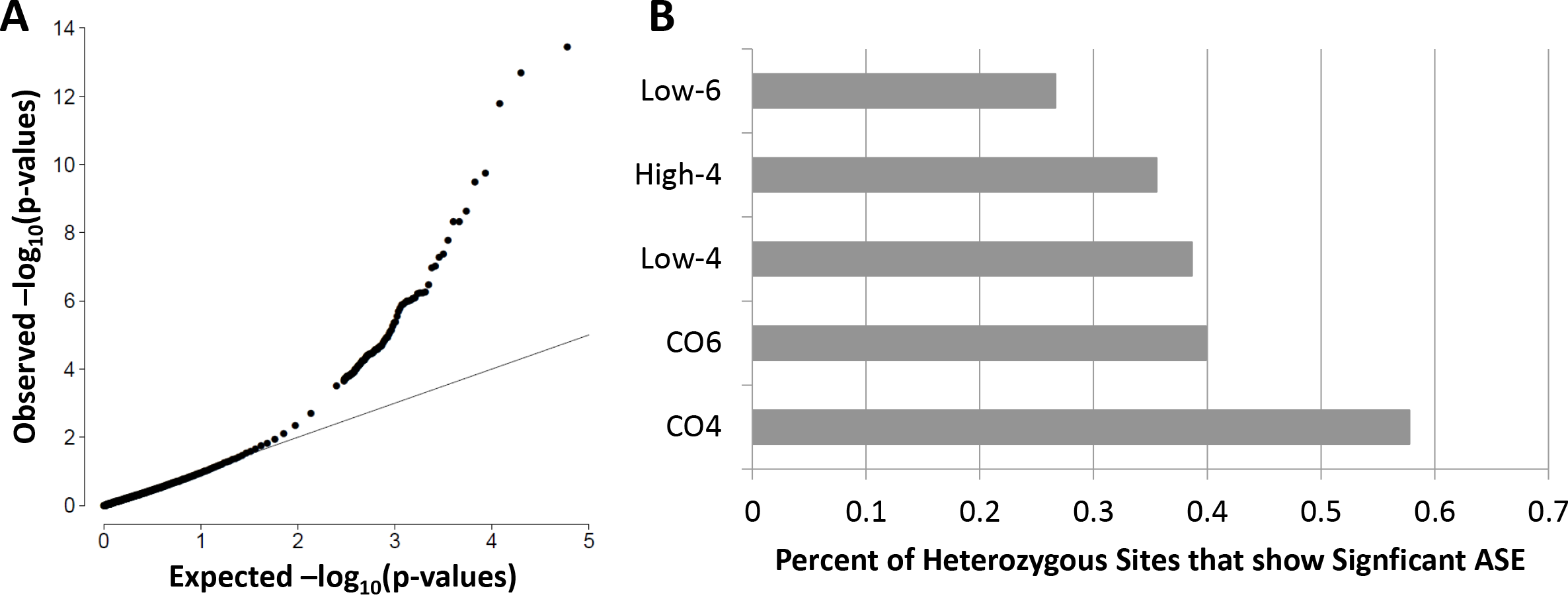
Allele-specific expression in colonocytes following co-culturing with microbiome. A) QQ-plot depicting the ASE nominal p-values for heterozygous SNPs in human colonocytes. B) Percentage of SNPs with allele-specific expression in each of the 5 samples (3 treatments, 2 controls) normalized by the number of heterozygous sites covered by at least 20 reads.

We then formally tested whether host transcriptional response may be modulated by an interaction between host genetics and the microbiome. Previous studies have examined gene-by-environment interaction in response to infection by searching for response expression quantitative trait loci (reQTLs), where the genetic effect on gene expression is only present under certain conditions (67–70). However, this type of study requires many individuals to gain enough statistical power. Instead, we searched for gene-by-environment interactions by examining ASE conditional on the exposure to the microbiome (conditional ASE, cASE). We identified 12 SNPs in 12 different genes that show cASE under any of the three treatment conditions (empi2rical FDR < 12%) (**Figure 5A-B**, Table S7 and Figure S3). These genes represent host response that is regulated by both host genetics and the interaction with the gut microbiome.

**Figure 5:**
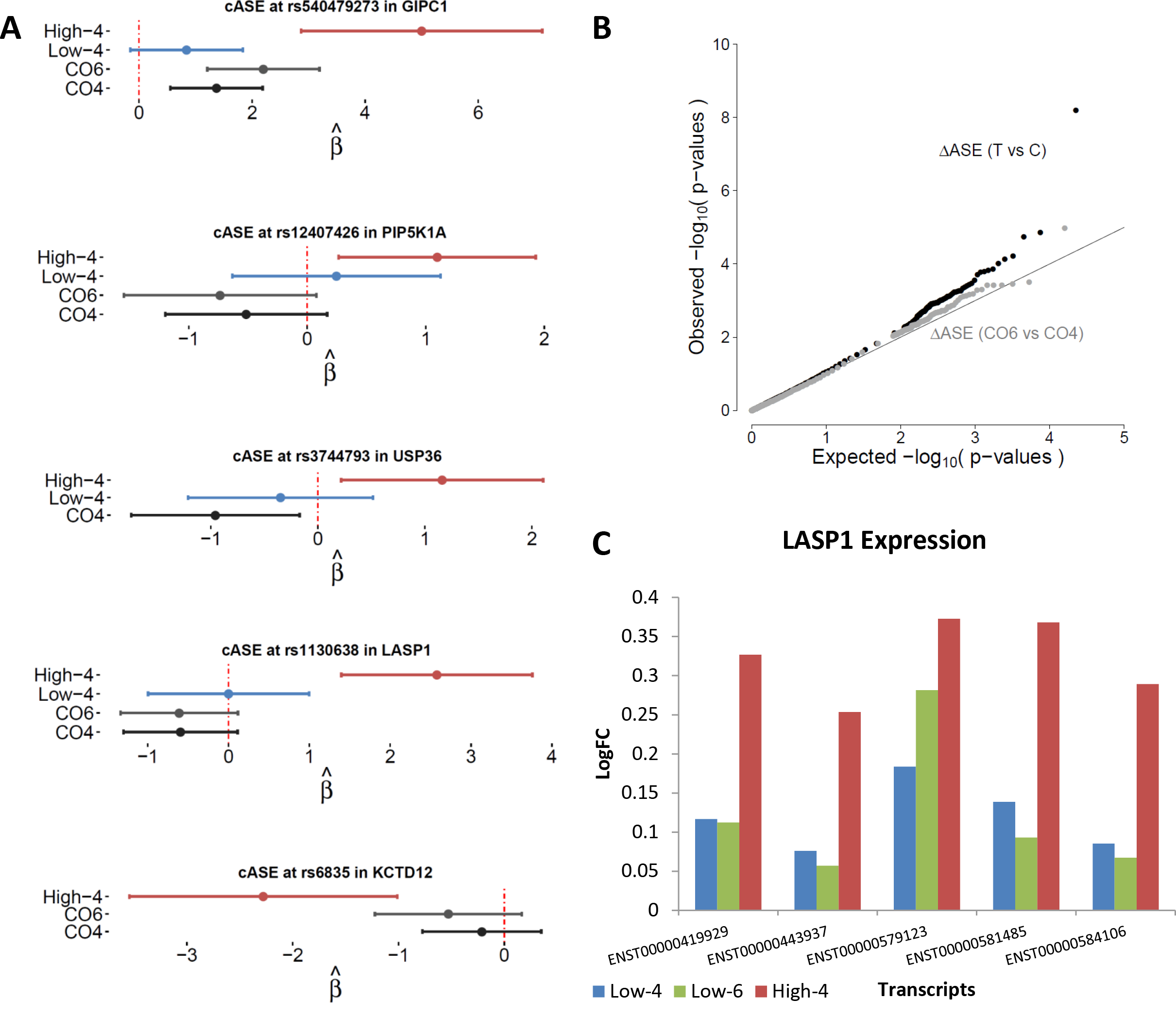
Gene-by-environment interaction in human colonocytes. A) Forest plots depicting conditional allele-specific expression (cASE) for 5 SNPs. Allele-specific expression is shown for samples with at least 20 reads covering the indicated SNP. Positive **β** indicates allele-specific expression favoring the reference allele B) QQ-plot showing the nominal p-values of SNPs that could be tested for cASE (20 reads covering SNP in both a treatment and the corresponding control or for both CO6 and CO4). C) Gene expression changes in each treatment (as compared to the corresponding control) for each of 5 transcripts of *LASP1* expressed in colonocytes.

Two of the 12 genes with cASE have been implicated in the immune response (*USP36*, *PIP5K1A*), while 8 of them have been linked to a disease affected by dysbiosis in the gut (*AFAP1L2*, *PIP5K1A*, *GIPC1*, *ASAP2*, *USP36*, *RNF213*, *KCTD12*, *LASP1*) (71–81). For example, we find cASE at SNP rs1130638 in *LASP1* as well as increased total expression of *LASP1* following exposure to the high concentration of microbiome at 4 hours (**Figure 5A** and **5C**). This suggests that the gut microbiome has a stronger effect on *LASP1* upregulation in the presence of a specific allelic variant. *LASP1* encodes a protein that binds to actin and regulates the cytoskeleton, and it has previously been shown to increase in expression following infection. Specifically, infection with hepatitis B virus X increased *LASP1* expression and led to cell migration (82). However, when *LASP1* expression was knocked-down following exposure to the virus, subsequent cell migration and movement was also reduced. Furthermore, colorectal cancer cells also show higher expression of *LASP1*, suggesting that *LASP1* plays a similar role in colonocytes. Together, these data suggest another mechanism by which exposure to the microbiome may lead to cell migration and perhaps carcinogenesis through influencing ASE and genotype-dependent expression of *LASP1*.

## Discussion

The gut microbiome has been shown to be complex and variable under physiological and pathological conditions. While studies of the microbiome have become more common, in humans, they have been mostly limited to identifying associations between microbial communities and host phenotypes. Here, we have developed a novel approach to directly investigate the transcriptional changes induced by live microbial communities on host colonic epithelial cells and how these changes are modulated by host genotype. The advantage of this method as compared to *in vivo* studies in mice is that it allows for high-throughput testing of multiple microbiome and host combinations with quick assessment of the interaction. Previous studies examining the host-microbiome interaction have studied germ-free mice exposed to the gut microbiome of humans (10, 46). While these studies have generated important insight on host-microbiome interactions, they have distinct caveats and limitations. First, while the environment of mice can be well-controlled, the interaction of mice and their microbiome may differ from the interaction of humans and their microbiome. Additionally, mice can be expensive to maintain and they have limited genetic variation, making it difficult to investigate a large number of genetic variants and identify loci involved in the host cellular response. Studies in humans, on the other hand, are able to use natural genetic variation to identify loci associated with microbiome composition, but it is difficult to control for the impact of other environmental factors. Our *in vitro* system allows for the study of interaction between many microbiomes and host cell cultures at a relatively much lower cost. Another advantage of this system is the ability to determine the changes in host cell response and microbiome composition over a time course, allowing us to gain insight on the cascade of transcriptional pathways involved in the response. Even though our system only focuses on one cell type and does not fully recapitulate the complexity of cell types and interactions in the gut, our data suggests that it provides a good representation of the results seen in *in vivo* studies in mice.

In addition to our novel experimental design, our analysis also adds to the understanding of the interaction between human genetic variation and the microbiome. Previous work has searched for quantitative trait loci that are associated with the abundance of certain bacteria but have lacked power to detect many loci (32, 33, 35). Our analysis of allele-specific expression maximizes the information available for each individual and allowed us to identify 12 loci that demonstrated conditional allele-specific expression and evidence of gene-by-microbiome interaction in a single individual. This system is easily amenable to scaling up in order to perform eQTL and response eQTL analysis (39, 63–66).

In this study we were able to learn about human colonocyte response to fecal microbial communities. We identified over 6,000 host genes that change expression following co-culture with the microbiome. These genes are enriched for certain functions including cell-cell interaction and cell migration, and in higher concentrations of microbiome, we see enrichment for genes involved in the immune response. When we further searched for genes where genetic variation affects the response to microbiome exposure, we found 12 genes containing cASE. Several of these genes can be linked to cell adhesion and migration (*AFAP1L2*, *PIP5K1A*, *GIPC1*, *ARFGAP3*, *ASAP2*) and *LASP1* has also been shown to change expression in colorectal cancer (80, 81, 83). These interactions demonstrate how the microbiome may influence cell-cell junctions and cell surface receptors, likely due to the *in vivo* reaction of colonocytes to protect the body from infection by sealing tight junctions and replacing cells that have been sloughed off by intestinal movement (48, 84–86). Among the genes with cASE, we also identified several genes that have been associated with diabetes (*GIPC1*, *USP36*, *RNF213*, *KCTD12*) or obesity (*PIP5K1A*) *(74, 75, 77–79)*. Both diabetes and obesity have been linked to microbiome composition (7, 27, 54). These genes may play a role in host-microbiome interactions and the dysbiosis that leads to these diseases.

Our study demonstrates a scalable approach to study host-gut microbiome interactions that depicts the *in vivo* relationship. This technique allowed us to start deciphering the impact of the microbiome on host cells and will help to determine how the microbiome may lead to disease through its influence on host cell gene regulation. We also highlight the importance of gene-by-microbiome interactions and suggest that it is not simply the genetics of an individual but the interplay between genetics and microbiome that will influence health and disease. Future studies using this approach with multiple individuals and microbiomes will identify key host factors and microbial communities that jointly influence human disease.

## Materials and methods

### Cell culture and treatment

Experiments were conducted using primary human colonic epithelial cells (HCoEpiC, lot: 9810), which we also term, colonocytes (ScienCell 2950). The cells were cultured on plates or flasks coated with poly-L-lysine (PLL) according to manufacturer’s specifications (ScienCell 0413). Colonocytes were cultured in colonic epithelial cell medium supplemented with colonic epithelial cell growth supplement and penicillin/streptomycin according to manufacturer's protocol (ScienCell 2951) at 37°C with 5% CO_2_. 24 hours before treatment, cells were changed to antibiotic-free media and moved to an incubator at 37°C, 5% CO_2_, and 5% O_2_.

Fecal extract was purchased from OpenBiome and arrived frozen on dry ice. Extract was not thawed until the day of treatment. Fecal extract was collected from a healthy, 22 year old male (Unit ID: 02-028-C). Prior to treatment, the fecal extract was thawed at 30°C and the microbial density was assessed by spectrophotometer (OD600) (Bio-Rad SmartSpec 3000). Media was removed from the colonocytes and fresh antibiotic-free media was added to the cells with a final microbial ratio of 10:1 or 100:1 microbe:colonocyte in each well (Low and High, respectively). Additional wells containing only colonocytes were also cultured in the same 24-well plate to be used as controls.

Following 4 or 6 hours, the wells were scraped on ice, pelleted and washed with cold PBS and then resuspended in lysis buffer (Dynabeads mRNA Direct Kit) and stored at −80°C until extraction of colonocyte RNA. Control treatments and Low-6 were done in triplicate while the Low-4 and High-4 were done in duplicate. The colonocytes exposed to the high concentration of microbiome for 6 hours were unhealthy and RNA was unable to be collected.

### RNA-library preparation from colonocytes

Poly-adenylated mRNAs were isolated from thawed cell lysates using the Dynabeads mRNA Direct Kit (Ambion) and following the manufacturer’s instructions. RNA-seq libraries were prepared using a protocol modified from the NEBNext Ultradirectional (NEB) library preparation protocol to use Barcodes from BIOOScientific added by ligation, as described in (87). The individual libraries were quantified using the KAPA real-time PCR system, following the manufacturer's instructions and using a custom-made series of standards obtained from serial dilutions of the phi-X DNA (Illumina). The libraries were then pooled and sequenced on two lanes of the Illumina Next-seq 500 in the Luca/Pique laboratory using the high output kits for 75 cycles and 300 cycles to obtain paired-end reads for an average of 150 million and 50 million total reads per sample, respectively.

### 16S rRNA gene sequencing and analysis of the microbiome preparation

Microbial DNA was extracted from the uncultured microbiome sample in triplicate using the PowerSoil kit from MO BIO Laboratories as directed, with a few modifications. Briefly, the fecal extract was spun to collect live microbes. The pellet was then resuspended in 200μL of phenol:chloroform and added to the 750μL bead solution from the PowerSoil kit. The kit protocol was then followed and the column was eluted in 60μL. This eluate was then purified using MinElute PCR Purification Kit (Qiagen) according to the manufacturer’s instructions.

16S rRNA gene amplification and sequencing was performed at the University of Minnesota Genomics Center (UMGC), as described in Burns et al. (12). Briefly, DNA isolated from the fecal extract was quantified by qPCR, and the V5-V6 regions of the 16S rRNA gene were PCR amplified. Nextera indexing primers were added in the first PCR using the V5F primer 5’-AATGATACGGCGACCACCGAGATCTACAC[i5]TCGTCGGCAGCGTC-3’, and V6R 5’-CAAGCAGAAGACGGCATACGAGAT[i7]GTCTCGTGGGCTCGG-3’, where [i5] and [i7] refer to the index sequences used by Illumina. This PCR was carried out using the KAPA HiFidelity Hot Start polymerase (Kapa Biosystems) for 20 cycles. The amplicons were then diluted 1:100 and used as input for a second PCR using different combinations of forward and reverse indexing primers for another 10 cycles. The pooled, size-selected product was diluted to 8pM, spiked with 15% PhiX and loaded onto an Illumina MiSeq instrument to generate the 16S rRNA gene sequences (v3 kit, PE 2 × 300), resulting in 2.2 million raw reads per sample, on average. Barcodes were removed from the sample reads by UMGC and the Nextera adaptors were trimmed using CutAdapt 1.8.1.

The trimmed 16S rRNA gene sequence pairs were quality filtered (q-score > 20, using QIIME 1.8.0) resulting in 1.41, 1.06, and 1.53 million high quality reads for sample replicates 1, 2, and 3, respectively (43, 44). OTUs were picked using the closed reference algorithm against the Greengenes database (August, 2013 release) (12, 43, 44, 88). The resulting OTU table was analyzed to determine microbial community diversity using QIIME scripts and rarefying to 280,000 reads.

### RNA sequencing and differential gene expression analysis

Reads were aligned to the hg19 human reference genome using STAR (89)(https://github.com/alexdobin/STAR/releases, version STAR_2.4.0h1), and the Ensemble reference transcriptome (version 75) with the following options:

STAR --runThreadN 12 --genomeDir <genome>

> --readFilesIn <fastqs.gz> --readFilesCommand zcat
>
> --outFileNamePrefix <stem> --outSAMtype BAM Unsorted
>
> --genomeLoad LoadAndKeep

where <genome> represents the location of the genome and index files, <fastqs.gz> represents that sample's fastq files, and <stem> represents the filename stem of that sample. For each sample, we merged sequencing replicates from the 2 different sequencing runs using samtools (version 2.25.0). We further required a quality score of 10 to remove reads mapping to multiple locations. We used the WASP suite of tools (vandeGeijn2015) (https://github.com/bmvdgeijn/WASP, downloaded 09/15/15) for allele-specific mapping and removing duplicates to ensure that there is no mapping bias at SNPs. The resulting alignments are used for the following analyses and the read counts can be found in Table S6.

To identify differentially expressed (DE) genes, we used DESeq2 (45) (R version 3.2.1, DESeq2 version 1.8.1) over experimental replicates for each treatment condition. DESeq2 was performed over each transcript expressed in all samples. A transcript was differentially expressed when the log2(fold-change) was greater than 0.25 and had a Benjamini-Hochberg adjusted p-value (90) < 0. 1. A gene was considered DE if at least one of its transcripts was DE.

### Gene ontology analysis

We utilized GeneTrail (91) to find enrichment of gene ontology terms. We compiled a list of unique genes that changed gene expression under any of the 3 conditions (Low-4, High-4, and Low-6) and determined which GO categories were under/over-represented as compared to a list of all genes expressed in colonocytes (15,781 genes). We considered a category over/under-represented if the Benjamini-Hochberg adjusted p-value < 0.05. Figure 3A depicts the top 10 categories over-represented that had an expected number of genes between 10 and 500. Enrichment is calculated by dividing the observed number of genes in a category by the expected number based on the total gene set.

### Enrichment of DE genes among genome-wide association studies

We downloaded the GWAS catalog (62)(version 1.0.1) on January 5th, 2016. To identify the overlap between DE genes in our dataset and those associated with a GWAS trait, we intersected genes that contain transcripts that change significantly and in the same direction in all 3 treatments with the reported genes from the GWAS catalog. We report enrichment with specific categories from the GWAS catalog: “Obesity-related traits”, “Inflammatory bowel disease”, “Ulcerative colitis”, “Colorectal cancer”, “Type 2 diabetes”, and “Crohn’s disease”. We used a Fisher's exact test on a 2×2 contingency table using 2 groups: genes that contain transcripts that are DE, in the same direction, in the 3 treatments (“ALL”) and other genes that are expressed in each sample (“NOT”). We then split these groups into 2: genes that are associated with the select disease (“TRAIT”) and genes that are associated with any other trait in the GWAS catalog (“OTHER GWAS”). Values are shown in Table S4.

### Joint genotyping and ASE inference

First, we identified SNPs to be studied for allele-specific expression (ASE). We used all 1KG SNPs from the phase 3 release (v5b.20130502, downloaded on 08/24/15) but removed SNPs if their minor allele frequency was less than 5% or they were found in annotated regions of copy number variation and ENCODE blacklisted regions (39). The resulting 7,340,521 SNPs were then studied in the following analysis.

Using samtools mpileup and the hg19 human reference genome, we obtained the read counts at each SNP in each sample from the RNA-seq data. These pileups were then processed using QuASAR package (66) by combining the RNA-seq reads from each sample (as they are all derived from the same colonocyte cell line) for joint genotyping. From the genotype information we identified heterozygous SNPs with read coverage of at least 20 and we tested them for ASE using QuASAR (66). Though we used combined read counts for genotyping, in order to identify ASE, we studied each sample separately but still combined the respective reads from the 75 and 300 cycle runs and across experimental replicates.

## Analysis of cASE

To identify conditional allele-specific expression (cASE) we transformed QuASAR β parameters to differential Z-scores (Z_Δ_) using the following formula:

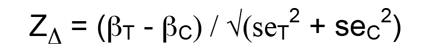

where β and se represent the estimates for the ASE parameter and its standard error for either the treatment (T) or control (C) samples.

The Z_Δ_ scores were then normalized by the standard deviation across Z_Δ_ scores corresponding to control versus control (controls at 4 and 6 hours).Finally p-values (p_Δ_) were calculated from the Z_Δ_ scores as p_Δ_ = 2 × pnorm(−|z|).

Under the null, Z_Δ_ are asymptotically normally distributed. To further correct for this small deviation we used the control versus control p-values to empirically estimate the FDR. The list of significant cASE SNPs (empirical FDR < 12%) is in Table S7.

## Accession numbers for sequencing data

Submission of 16S sequencing data of uncultured microbiome and RNA sequencing data of colonocytes in all conditions to Short Read Archive (SRA) is pending.

## Acknowledgements

We thank Emily Davenport, Sonia Kupfer and members of the Luca, Blekhman and Pique-Regi groups for helpful comments. We thank the Wayne State University High Performance Computing Grid and the Minnesota Supercomputing Institute for computational support. We thank OpenBiome for supplying the fecal extract for treatment.

## Supplemental Figure and Table Legends

**Figure S1**: Volcano plots of host gene expression changes following exposure to the microbiome. Each plot depicts gene expression changes for a different treatment as compared to the time-specific control sample. X-axis shows the log_2_FC and the y-axis shows the −log_10_(Benjamini-Hochberg adjusted p-value). The points colored light blue are significantly differentially expressed (Benjamini-Hochberg adjusted p-value < 0.1, |logFC| > 0.25).

**Figure S2**:: Functional categories of DE genes at High-4. Enrichment of immune-related GO categories of genes that are significantly increased in expression in High-4 as compared to Low-4.

**Figure S3**: Examples of cASE following exposure to the microbiome. Forest plots depicting conditional allele-specific expression (cASE) for 7 SNPs. ASE is shown for samples where at least 20 reads cover the indicated SNP (positive β indicates ASE favoring the reference allele).

**Table S1**: 16S rRNA gene sequencing analysis of microbiome composition. Sequencing of fecal extract was done in triplicate. The table shows proportional data.

**Table S2**: Differentially expressed genes in colonocytes following co-culturing. Low-4 and High-4 are compared to CO4 while Low-6 is compared to CO6 to determine gene expression changes. Changes are shown for each transcript with the gene IDs and symbols in the last 2 columns.

**Table S3**: Differences in gene expression between High-4 and Low-4. Changes in expression are shown for each transcript.

**Table S4**: Enrichment of GWAS traits among DE genes. Six traits that have previously been linked to the microbiome were assessed. A 2×2 contingency table was constructed: “ALL” (genes that contain transcripts that are differentially expressed, in the same direction, in all three treatments) or “NOT” (any other gene expressed in this study), “TRAIT” (genes that are associated with the shown trait in the GWAS catalog) or “OTHER GWAS” (genes associated with another trait in the GWAS catalog). A Fisher’s Exact test was performed to obtain the p-values and odds ratios.

**Table S5**: Allele-specific expression in colonocytes. ASE is shown for each SNP in the treatment or control sample in which ASE is found. Positive β indicates higher expression for the reference allele.

**Table S6**: Allele-specific expression as a function of sequencing depth.

**Table S7**: Conditional allele-specific expression following exposure to microbiome. ASE is shown for the treatment in which cASE occurs and its respective control. dASE values indicate the significant difference between ASE in the treatment and control.

## References

1. Lozupone CA, Stombaugh JI, Gordon JI, Jansson JK, Knight R. 2012. Diversity, stability and resilience of the human gut microbiota. Nature 489:220–230.

2. Gevers D, Kugathasan S, Denson LA, Vázquez-Baeza Y, Van Treuren W, Ren B, Schwager E, Knights D, Song SJ, Yassour M, Morgan XC, Kostic AD, Luo C, González A, McDonald D, Haberman Y, Walters T, Baker S, Rosh J, Stephens M, Heyman M, Markowitz J, Baldassano R, Griffiths A, Sylvester F, Mack D, Kim S, Crandall W, Hyams J, Huttenhower C, Knight R, Xavier RJ. 2014. The treatment-naive microbiome in new-onset Crohn's disease. Cell Host Microbe 15:382–392.

3. Larsen N, Vogensen FK, van den Berg FWJ, Nielsen DS, Andreasen AS, Pedersen BK, Al-Soud WA, Sørensen SJ, Hansen LH, Jakobsen M. 2010.Gut microbiota in human adults with type 2 diabetes differs from nondiabetic adults. PLoS One 5:e9085.

4. Zhang X, Shen D, Fang Z, Jie Z, Qiu X, Zhang C, Chen Y, Ji L. 2013. Human gut microbiota changes reveal the progression of glucose intolerance. PLoS One 8:e71108.

5. Lin HV, Frassetto A, Kowalik EJ Jr, Nawrocki AR, Lu MM, Kosinski JR, Hubert JA, Szeto D, Yao X, Forrest G, Marsh DJ. 2012. Butyrate and propionate protect against diet-induced obesity and regulate gut hormones via free fatty acid receptor 3-independent mechanisms. PLoS One 7:e35240.

6. Turnbaugh PJ, Ley RE, Mahowald MA, Magrini V, Mardis ER, Gordon JI. 2006.An obesity-associated gut microbiome with increased capacity for energy harvest. Nature 444:1027–1031.

7. Ley RE, Bäckhed F, Turnbaugh P, Lozupone CA, Knight RD, Gordon JI. 2005.Obesity alters gut microbial ecology. Proc Natl Acad Sci U S A 102:11070–11075.

8. Schwiertz A, Taras D, Schäfer K, Beijer S, Bos NA, Donus C, Hardt PD. 2010. Microbiota and SCFA in lean and overweight healthy subjects. Obesity 18:190–195.

9. Turnbaugh PJ, Bäckhed F, Fulton L, Gordon JI. 2008. Diet-induced obesity is linked to marked but reversible alterations in the mouse distal gut microbiome. Cell Host Microbe 3:213–223.

10. Goodrich JK, Waters JL, Poole AC, Sutter JL, Koren O, Blekhman R, Beaumont M, Van Treuren W, Knight R, Bell JT, Spector TD, Clark AG Ley RE. 2014. Human genetics shape the gut microbiome. Cell 159:789–799.

11. Sinha R, Ahn J, Sampson JN, Shi J, Yu G, Xiong X, Hayes RB, Goedert JJ. 2016. Fecal Microbiota,Fecal Metabolome, and Colorectal Cancer Interrelations. PLoS One 11:e0152126.

12. Burns MB, Lynch J, Starr TK, Knights D, Blekhman R. 2015. Virulence genes are a signature of the microbiome in the colorectal tumor microenvironment. Genome Med 7:55.

13. Liu X, Zou Q, Zeng B, Fang Y, Wei H. 2013. Analysis of fecal Lactobacillus community structure in patients with early rheumatoid arthritis. Curr Microbiol 67:170–176.

14. Vaahtovuo J, Munukka E, Korkeamäki M, Luukkainen R, Toivanen P. 2008.Fecal microbiota in early rheumatoid arthritis. J Rheumatol 35:1500–1505.

15. Scheperjans F, Aho V, Pereira PAB, Koskinen K, Paulin L, Pekkonen E, Haapaniemi E, Kaakkola S, Eerola-Rautio J, Pohja M, Kinnunen E, Murros K, Auvinen P. 2015. Gut microbiota are related to Parkinson’s disease and clinical phenotype. Mov Disord 30:350–358.

16. Mulak A, Bonaz B. 2015. Brain-gut-microbiota axis in Parkinson’s disease. World J Gastroenterol 21:10609–10620.

17. Qin J, Li R, Raes J, Arumugam M, Burgdorf KS, Manichanh C, Nielsen T, Pons N, Levenez F, Yamada T, Mende DR, Li J, Xu J, Li S, Li D, Cao J, Wang B, Liang H, Zheng H, Xie Y, Tap J, Lepage P, Bertalan M, Batto J-M, Hansen T, Le Paslier D, Linneberg A, Nielsen HB, Pelletier E, Renault P, Sicheritz-Ponten T, Turner K, Zhu H, Yu C, Li S, Jian M, Zhou Y, Li Y, Zhang X, Li S, Qin N, Yang H, Wang J, Brunak S, Doré J, Guarner F, Kristiansen K, Pedersen O, Parkhill J, Weissenbach J, MetaHIT Consortium, Bork P, Ehrlich SD, Wang J. 2010. A human gut microbial gene catalogue established by metagenomic sequencing. Nature 464:59–65.

18. Human Microbiome Project Consortium. 2012. Structure, function and diversity of the healthy human microbiome. Nature 486:207–214.

19. Praveen P, Jordan F, Priami C, Morine MJ. 2015. The role of breastfeeding in infant immune system: a systems perspective on the intestinal microbiome. Microbiome 3:41.

20. Bezirtzoglou E, Tsiotsias A, Welling GW. 2011. profile in feces of breast-and formula-fed newborns by using fluorescence in situ hybridization (FISH). Anaerobe 17:478–482.

21. Penders J, Thijs C, Vink C, Stelma FF, Snijders B, Kummeling I, van den Brandt PA, Stobberingh EE. 2006. Factors influencing the composition of the intestinal microbiota in early infancy. Pediatrics 118:511–521.

22. Dominianni C, Sinha R, Goedert JJ, Pei Z, Yang L, Hayes RB, Ahn J. 2015.Sex, body mass index, and dietary fiber intake influence the human gut microbiome. PLoS One 10:e0124599.

23. Bolnick DI, Snowberg LK, Hirsch PE, Lauber CL, Org E, Parks B, Lusis AJ, Knight R, Caporaso JG, Svanbäck R. 2014. Individual diet has sex-dependent effects on vertebrate gut microbiota. Nat Commun 5:4500.

24. Turnbaugh PJ, Ridaura VK, Faith JJ, Rey FE, Knight R, Gordon JI. 2009. The effect of diet on the human gut microbiome: a metagenomic analysis in humanized gnotobiotic mice. Sci Transl Med 1:6ra14.

25. Lee S, Sung J, Lee J, Ko G. 2011. Comparison of the gut microbiotas of healthy adult twins living in South Korea and the United States. Appl Environ Microbiol 77:7433–7437.

26. Tims S, Derom C, Jonkers DM, Vlietinck R, Saris WH, Kleerebezem M, de Vos WM, Zoetendal EG. 2013. Microbiota conservation and BMI signatures in adult monozygotic twins. ISME J 7:707–717.

27. Turnbaugh PJ, Hamady M, Yatsunenko T, Cantarel BL, Duncan A, Ley RE, Sogin ML, Jones WJ, Roe BA, Affourtit JP, Egholm M, Henrissat B, Heath AC, Knight R, Gordon JI. 2009. A core gut microbiome in obese and lean twins. Nature 457:480–484.

28. Yatsunenko T, Rey FE, Manary MJ, Trehan I, Dominguez-Bello MG, Contreras M, Magris M, Hidalgo G, Baldassano RN, Anokhin AP, Heath AC, Warner B, Reeder J, Kuczynski J, Caporaso JG, Lozupone CA, Lauber C, Clemente JC, Knights D, Knight R, Gordon JI. 2012. Human gut microbiome viewed across age and geography. Nature 486:222–227.

29. Benson AK, Kelly SA, Legge R, Ma F, Low SJ, Kim J, Zhang M, Oh PL, Nehrenberg D, Hua K, Kachman SD, Moriyama EN, Walter J, Peterson DA, Pomp D. 2010. Individuality in gut microbiota composition is a complex polygenic trait shaped by multiple environmental and host genetic factors. Proc Natl Acad Sci U S A 107:18933–18938.

30. McKnite AM, Perez-Munoz ME, Lu L, Williams EG, Brewer S, Andreux PA, Bastiaansen JWM, Wang X, Kachman SD, Auwerx J, Williams RW, Benson AK, Peterson DA, Ciobanu DC. 2012. Murine gut microbiota is defined by host genetics and modulates variation of metabolic traits. PLoS One 7:e39191.

31. Leamy LJ, Kelly SA, Nietfeldt J, Legge RM, Ma F, Hua K, Sinha R, Peterson DA, Walter J, Benson AK, Pomp D. 2014. Host genetics and diet, but not immunoglobulin A expression, converge to shape compositional features of the gut microbiome in an advanced intercross population of mice. Genome Biol 15:552.

32. Davenport ER, Cusanovich DA, Michelini K, Barreiro LB, Ober C, Gilad Y. 2015. Genome-Wide Association Studies of the Human Gut Microbiota. PLoS One 10:e0140301.

33. Blekhman R, Goodrich JK, Huang K, Sun Q, Bukowski R, Bell JT, Spector TD, Keinan A, Ley RE, Gevers D, Clark AG. 2015. Host genetic variation impacts microbiome composition across human body sites. Genome Biol 16:191.

34. Org E, Parks BW, Joo JWJ, Emert B, Schwartzman W, Kang EY, Mehrabian M, Pan C, Knight R, Gunsalus R, Drake TA, Eskin E, Lusis AJ. 2015. Genetic and environmental control of host-gut microbiota interactions. Genome Res 25:1558–1569.

35. Goodrich JK, Davenport ER, Waters JL, Clark AG, Ley RE. 2016. Crossspecies comparisons of host genetic associations with the microbiome. Science 352:532–535.

36. GTEx Consortium. 2013. The Genotype-Tissue Expression (GTEx) project. Nat Genet 45:580–585.

37. Stranger BE, Forrest MS, Dunning M, Ingle CE, Beazley C, Thorne N, Redon R, Bird CP, de Grassi A, Lee C, Tyler-Smith C, Carter N, Scherer SW, Tavare S, Deloukas P, Hurles ME, Dermitzakis ET. 2007. Relative impact of nucleotide and copy number variation on gene expression phenotypes. Science 315:848–853.

38. Kasowski M, Grubert F, Heffelfinger C, Hariharan M, Asabere A, Waszak SM, Habegger L, Rozowsky J, Shi M, Urban AE, Hong M-Y, Karczewski KJ, Huber W, Weissman SM, Gerstein MB, Korbel JO, Snyder M. 2010. Variation in transcription factor binding among humans. Science 328:232–235.

39. Degner JF, Pai AA, Pique-Regi R, Veyrieras J-B, Gaffney DJ, Pickrell JK, De Leon S, Michelini K, Lewellen N, Crawford GE, Others. 2012. DNase I sensitivity QTLs are a major determinant of human expression variation. Nature 482:390–394.

40. Gibbs JR, van der Brug MP, Hernandez DG, Traynor BJ, Nalls MA, Lai SL, Arepalli S, Dillman A, Rafferty IP, Troncoso J, Johnson R, Zielke HR, Ferrucci L, Longo DL, Cookson MR, Singleton AB. 2010. Abundant quantitative trait loci exist for DNA methylation and gene expression in human brain. PLoS Genet 6:e1000952.

41. Pastinen T. 2010. Genome-wide allele-specific analysis: insights into regulatory variation. Nat Rev Genet 11:533–538.

42. McDaniell R, Lee B-K, Song L, Liu Z, Boyle AP, Erdos MR, Scott LJ, Morken MA, Kucera KS, Battenhouse A, Keefe D, Collins FS, Willard HF, Lieb JD, Furey TS, Crawford GE, Iyer VR, Birney E. 2010. Heritable individual-specific and allele-specific chromatin signatures in humans. Science 328:235–239.

43. Navas-Molina JA, Peralta-Sanchez JM, Gonzalez A, McMurdie PJ, Vazquez-Baeza Y, Xu Z, Ursell LK, Lauber C, Zhou H, Song SJ, Huntley J, Ackermann GL, Berg-Lyons D, Holmes S, Caporaso JG, Knight R. 2013. Advancing our understanding of the human microbiome using QIIME. Methods Enzymol 531:371–444.

44. Caporaso JG, Kuczynski J, Stombaugh J, Bittinger K, Bushman FD, Costello EK, Fierer N, Peña AG, Goodrich JK, Gordon JI, Huttley GA, Kelley ST, Knights D, Koenig JE, Ley RE, Lozupone CA, McDonald D, Muegge BD, Pirrung M, Reeder J, Sevinsky JR, Turnbaugh PJ, Walters WA, Widmann J, Yatsunenko T, Zaneveld J, Knight R. 2010. QIIME allows analysis of high-throughput community sequencing data. Nat Methods 7: 335–336.

45. Love MI, Huber W, Anders S. 2014. Moderated estimation of fold change and dispersion for RNA-seq data with DESeq2. Genome Biol 15:550.

46. Camp JG, Frank CL, Lickwar CR, Guturu H, Rube T, Wenger AM, Chen J, Bejerano G, Crawford GE, Rawls JF. 2014. Microbiota modulate transcription in the intestinal epithelium without remodeling the accessible chromatin landscape. Genome Res 24:1504–1516.

47. Fukushima K, Ogawa H, Takahashi K, Naito H, Funayama Y, Kitayama T, Yonezawa H, Sasaki I. 2003. Non-pathogenic bacteria modulate colonic epithelial gene expression in germ-free mice. Scand J Gastroenterol 38:626–634.

48. Ewaschuk JB, Diaz H, Meddings L, Diederichs B, Dmytrash A, Backer J, Looijer-van Langen M, Madsen KL. 2008. Secreted bioactive factors from Bifidobacterium infantis enhance epithelial cell barrier function. Am J Physiol Gastrointest Liver Physiol 295:G1025–34.

49. Larsson E, Tremaroli V, Lee YS, Koren O, Nookaew I, Fricker A, Nielsen J, Ley RE, Bäckhed F. 2012. Analysis of gut microbial regulation of host gene expression along the length of the gut and regulation of gut microbial ecology through MyD88. Gut 61:1124–1131.

50. Moon Y, Yang H, Kim YB. 2007. Up-regulation of early growth response gene 1 (EGR-1) via ERK1/2 signals attenuates sulindac sulfide-mediated cytotoxicity in the human intestinal epithelial cells. Toxicol Appl Pharmacol 223:155–163.

51. Chen G, Li H, Niu X, Li G, Han N, Li X, Li G, Liu Y, Sun G, Wang Y, Li Z, Li Q. 2015. Identification of key genes associated with colorectal cancer based on the transcriptional network. Pathol Oncol Res 21:719–725.

52. El Aidy S, van Baarlen P, Derrien M, Lindenbergh-Kortleve DJ, Hooiveld G, Levenez F, Doré J, Dekker J, Samsom JN, Nieuwenhuis EES, Kleerebezem M. 2012. Temporal and spatial interplay of microbiota and intestinal mucosa drive establishment of immune homeostasis in conventionalized mice. Mucosal Immunol 5:567–579.

53. Rawls JF, Mahowald MA, Ley RE, Gordon JI. 2006. Reciprocal gut microbiota transplants from zebrafish and mice to germ-free recipients reveal host habitat selection. Cell 127:423–433.

54. Qin J, Li Y, Cai Z, Li S, Zhu J, Zhang F, Liang S, Zhang W, Guan Y, Shen D, Peng Y, Zhang D, Jie Z, Wu W, Qin Y, Xue W, Li J, Han L, Lu D, Wu P, Dai Y, Sun X, Li Z, Tang A, Zhong S, Li X, Chen W, Xu R, Wang M, Feng Q, Gong M, Yu J, Zhang Y, Zhang M, Hansen T, Sanchez G, Raes J, Falony G, Okuda S, Almeida M, LeChatelier E, Renault P, Pons N, Batto J-M, Zhang Z, Chen H, Yang R, Zheng W, Li S, Yang H, Wang J, Ehrlich SD, Nielsen R, Pedersen O, Kristiansen K, Wang J. 2012. A metagenome-wide association study of gut microbiota in type 2 diabetes. Nature 490:55–60.

55. He Q, Li X, Liu C, Su L, Xia Z, Li X, Li Y, Li L, Yan T, Feng Q, Xiao L. 2016.Dysbiosis of the fecal microbiota in the TNBS-induced Crohn’s disease mouse model. Appl Microbiol Biotechnol 100:4485–4494.

56. Hoffmann TW, Pham H-P, Bridonneau C, Aubry C, Lamas B, Martin-Gallausiaux C, Moroldo M, Rainteau D, Lapaque N, Six A, Richard ML, Fargier E, Le Guern M-E, Langella P, Sokol H. 2016. Microorganisms linked to inflammatory bowel disease-associated dysbiosis differentially impact host physiology in gnotobiotic mice. ISME J 10:460–477.

57. Walujkar SA, Dhotre DP, Marathe NP, Lawate PS, Bharadwaj RS, Shouche YS. 2014. Characterization of bacterial community shift in human Ulcerative Colitis patients revealed by Illumina based 16S rRNA gene amplicon sequencing. Gut Pathog 6:22.

58. Vannucci L, Stepankova R, Kozakova H, Fiserova A, Rossmann P, Tlaskalova-Hogenova H. 2008. Colorectal carcinogenesis in germ-free and conventionally reared rats: different intestinal environments affect the systemic immunity. Int J Oncol 32:609–617.

59. Li Y, Kundu P, Seow SW, de Matos CT, Aronsson L, Chin KC, Karre K, Pettersson S, Greicius G. 2012. Gut microbiota accelerate tumor growth via c-jun and STAT3 phosphorylation in APCMin/+ mice. Carcinogenesis 33:1231–1238.

60. Gagniere J, Raisch J, Veziant J, Barnich N, Bonnet R, Buc E, Bringer M-A, Pezet D, Bonnet M. 2016. Gut microbiota imbalance and colorectal cancer. World J Gastroenterol 22:501–518.

61. Marchesi JR, Dutilh BE, Hall N, Peters WHM, Roelofs R, Boleij A, Tjalsma H. 2011. Towards the human colorectal cancer microbiome. PLoS One 6:e20447a.

62. Welter D, MacArthur J, Morales J, Burdett T, Hall P, Junkins H, Klemm A, Flicek P, Manolio T, Hindorff L, Parkinson H. 2014. The NHGRI GWAS Catalog, a curated resource of SNP-trait associations. Nucleic Acids Res 42:D1001–6.

63. Salava A, Lauerma A. 2014. Role of the skin microbiome in atopic dermatitis. Clin Transl Allergy 4:33.

64. Sellitto M, Bai G, Serena G, Fricke WF, Sturgeon C, Gajer P, White JR, Koenig SSK, Sakamoto J, Boothe D, Gicquelais R, Kryszak D, Puppa E, Catassi C, Ravel J, Fasano A. 2012. Proof of concept of microbiome-metabolome analysis and delayed gluten exposure on celiac disease autoimmunity in genetically at-risk infants. PLoS One 7:e33387.

65. Kostic AD, Xavier RJ, Gevers D. 2014. The microbiome in inflammatory bowel disease: current status and the future ahead. Gastroenterology 146:1489–1499.

66. Harvey CT, Moyerbrailean GA, Davis GO, Wen X, Luca F, Pique-Regi R. 2015.QuASAR: quantitative allele-specific analysis of reads. Bioinformatics 31:1235–1242.

67. Siddle KJ, Deschamps M, Tailleux L, Nédélec Y, Pothlichet J, Lugo-Villarino G, Libri V, Gicquel B, Neyrolles O, Laval G, Patin E, Barreiro LB, Quintana-Murci L. 2014. A genomic portrait of the genetic architecture and regulatory impact of microRNA expression in response to infection. Genome Res 24:850–859.

68. Caliskan M, Baker SW, Gilad Y, Ober C. 2015. Host genetic variation influences gene expression response to rhinovirus infection. PLoS Genet 11:e1005111.

69. Idaghdour Y, Quinlan J, Goulet J-P, Berghout J, Gbeha E, Bruat V, de Malliard T, Grenier J-C, Gomez S, Gros P, Rahimy MC, Sanni A, Awadalla P. 2012. Evidence for additive and interaction effects of host genotype and infection in malaria. Proc Natl Acad Sci U S A 109:16786–16793.

70. Barreiro LB, Tailleux L, Pai AA, Gicquel B, Marioni JC, Gilad Y. 2012. Deciphering the genetic architecture of variation in the immune response to Mycobacterium tuberculosis infection. Proceedings of the National Academy of Sciences 109:1204–1209.

71. Thevenon D, Engel E, Avet-Rochex A, Gottar M, Bergeret E, Tricoire H, Benaud C, Baudier J, Taillebourg E, Fauvarque M-O. 2009. The Drosophila ubiquitin-specific protease dUSP36/Scny targets IMD to prevent constitutive immune signaling. Cell Host Microbe 6:309–320.

72. Lee SY, Kim B, Yoon S, Kim YJ, Liu T, Woo JH, Chwae Y-J, Joe E-H, Jou I. 2010. Phosphatidylinositol 4-phosphate 5-kinase alpha is induced in ganglioside-stimulated brain astrocytes and contributes to inflammatory responses. Exp Mol Med 42:662–673.

73. Emaduddin M, Edelmann MJ, Kessler BM, Feller SM. 2008. Odin (ANKS1A) is a Src family kinase target in colorectal cancer cells. Cell Commun Signal 6:7.

74. Doumatey AP, Xu H, Huang H, Trivedi NS, Lei L, Elkahloun A, Adeyemo A, Rotimi CN. 2015. Global Gene Expression Profiling in Omental Adipose Tissue of Morbidly Obese Diabetic African Americans. J Endocrinol Metab 5:199–210.

75. Nguyen T-P, Liu W-C, Jordán F. 2011. Inferring pleiotropy by network analysis: linked diseases in the human PPI network. BMC Syst Biol 5:179.

76. Macadam RC, Sarela AI, Farmery SM, Robinson PA, Markham AF, Guillou PJ. 2000. Death from early colorectal cancer is predicted by the presence of transcripts of the REG gene family. Br J Cancer 83:188–195.

77. Greenawalt DM, Sieberts SK, Cornelis MC, Girman CJ, Zhong H, Yang X, Guinney J, Qi L, Hu FB. 2012. Integrating genetic association, genetics of gene expression, and single nucleotide polymorphism set analysis to identify susceptibility Loci for type 2 diabetes mellitus. Am J Epidemiol 176:423–430.

78. Cauchi S, Proenga C, Choquet H, Gaget S, De Graeve F, Marre M, Balkau B, Tichet J, Meyre D, Vaxillaire M, Froguel P, D.E.S.I.R. Study Group. 2008. Analysis of novel risk loci for type 2 diabetes in a general French population: the D.E.S.I.R. study. J Mol Med 86:341–348.

79. Kobayashi H, Yamazaki S, Takashima S, Liu W, Okuda H, Yan J, Fujii Y, Hitomi T, Harada KH, Habu T, Koizumi A. 2013. Ablation of Rnf213 retards progression of diabetes in the Akita mouse. Biochem Biophys Res Commun 432:519–525.

80. Zhao L, Wang H, Sun X, Ding Y. 2010. Comparative proteomic analysis identifies proteins associated with the development and progression of colorectal carcinoma. FEBS J 277:4195–4204.

81. Zhao L, Wang H, Liu C, Liu Y, Wang X, Wang S, Sun X, Li J, Deng Y,Jiang Y, Ding Y. 2010. Promotion of colorectal cancer growth and metastasis by the LIM and SH3 domain protein 1. Gut 59:1226–1235.

82. Tang R, Kong F, Hu L, You H, Zhang P, Du W, Zheng K. 2012. Role of hepatitis B virus X protein in regulating LIM and SH3 protein 1 (LASP-1) expression to mediate proliferation and migration of hepatoma cells. Virol J 9:163.

83. Wang H, Shi J, Luo Y, Liao Q, Niu Y, Zhang F, Shao Z, Ding Y, Zhao L. 2014. LIM and SH3 protein 1 induces TGFβ-mediated epithelial-mesenchymal transition in human colorectal cancer by regulating S100A4 expression. Clin Cancer Res 20:5835–5847.

84. Resta-Lenert S, Barrett KE. 2003. Live probiotics protect intestinal epithelial cells from the effects of infection with enteroinvasive Escherichia coli (EIEC). Gut 52:988–997.

85. Shimada Y, Kinoshita M, Harada K, Mizutani M, Masahata K, Kayama H, Takeda K. 2013. Commensal bacteria-dependent indole production enhances epithelial barrier function in the colon. PLoS One 8:e80604.

86. Cherbuy C, Honvo-Houeto E, Bruneau A, Bridonneau C, Mayeur C, Duée P-H, Langella P, Thomas M. 2010. Microbiota matures colonic epithelium through a coordinated induction of cell cycle-related proteins in gnotobiotic rat. Am J Physiol Gastrointest Liver Physiol 299:G348–57.

87. Moyerbrailean GA, Davis GO, Harvey CT, Watza D, Wen X, Pique-Regi R, Luca F. 2015. A high-throughput RNA-seq approach to profile transcriptional responses. bioRxiv.

88. DeSantis TZ, Hugenholtz P, Larsen N, Rojas M, Brodie EL, Keller K, Huber T, Dalevi D, Hu P, Andersen GL. 2006. Greengenes, a chimera-checked 16S rRNA gene database and workbench compatible with ARB. Appl Environ Microbiol 72:5069–5072.

89. Dobin A, Davis CA, Schlesinger F, Drenkow J, Zaleski C, Jha S, Batut P, Chaisson M, Gingeras TR. 2013. STAR: ultrafast universal RNA-seq aligner. Bioinformatics 29:15–21.

90. Benjamini Y, Hochberg Y. 1995. Controlling the False Discovery Rate: A Practical and Powerful Approach to Multiple Testing. J R Stat Soc Series B Stat Methodol 57:289–300.

91. Backes C, Keller A, Kuentzer J, Kneissl B, Comtesse N, Elnakady YA, Muller R, Meese E, Lenhof H-P. 2007. GeneTrail--advanced gene set enrichment analysis. Nucleic Acids Res 35:W186–92.

